# Out with the Old: Contrasting Histone Marks are Associated with Dosage Compensation on the Ancient and New Z of a Moth with Complex Sex Chromosomes

**DOI:** 10.64898/2026.02.02.702263

**Authors:** Vincent Kiplangat Bett, Ariana Macon, Marwan Elkrewi, Beatriz Vicoso

## Abstract

Sex chromosome differentiation is often accompanied by the evolution of dosage compensation (DC) mechanisms that balance gene expression between autosomes and sex chromosomes, and consequently between the sexes. ZW systems were traditionally thought to exhibit only partial DC, but recent studies in Lepidoptera, *Artemia franciscana*, and *Apalone spinifera* suggest a diverse range of compensation strategies, challenging traditional assumptions. While DC often involves chromatin-level regulation, the specific mechanisms in most ZW systems are still poorly understood. To explore these gaps, we generated a genome assembly of *Cameraria ohridella* (horse-chestnut leaf miner moth, Gracillidae), and combined transcriptomic data from two tissues with CUT&Tag epigenomic profiling targeting both active (H4K16ac, H3K4me3) and repressive (H3K27me3) chromatin marks. Our findings reveal a highly dynamic landscape of sex chromosome evolution in *C. ohridella*, including the ancestral Z (AncZ), a NeoZ_1_ formed by fusion of AncZ with an autosome, and a NeoZ_2_ that likely arose via the fusion of another autosome to the W chromosome. We uncover distinct dosage compensation (DC) patterns across ancestral and neo-sex chromosome regions. On the Ancestral Z, DC involves repression of gene expression in males (ZZ), with a depletion of the active histone mark H4K16ac being observed in this region. In contrast, NeoZ_1_ regions show upregulation of gene expression in the heterogametic sex (ZW), accompanied by enrichment of H4K16ac. Together, our findings underscore the dynamic nature of sex chromosome evolution and reveal variation in dosage compensation strategies across the Z chromosome, where the NeoZ_1_ appears to evolve entirely new, Drosophila-like mechanisms rather than co-opting the existing Nematode-like mechanisms of AncZ. We performed simulations that suggest that cooption of ancient DC mechanisms on a neo-sex chromosome is expected when it acts through up-regulation in the heterogametic sex but not when it involves downregulation in the homogametic sex, providing a framework for understanding the different patterns observed between Drosophila and Lepidoptera neo sex chromosomes.

## Introduction

Sex chromosomes have evolved independently multiple times in both animals and plants, often as XY (male heterogamety) or ZW (female heterogamety) systems (Graves 2008). They are acquired from pairs of initially homologous autosomes after the emergence of one or several sex-determining factor(s), typically followed by the evolution of recombination suppression between newly formed proto-X and Y (or Z and W) chromosomes. The absence of recombination results in progressive genetic degeneration of the Y chromosome (or W), characterized by accumulation of deleterious mutations and repeats, and by gene loss (Bachtrog 2013). The fusion of sex chromosomes with autosomes to form neo-sex chromosomes was generally considered a rare evolutionary event in systems with female heterogamety (Pokorná et al. 2014), and in particular species with high chromosome number such as Lepidoptera (where ZW females typically have 31 chromosomes) (Anderson et al. 2020). Despite this expectation, emerging evidence points to a surprising number of neo-sex chromosome formations within Lepidoptera (Wright et al. 2024; Fraïsse et al. 2017; Mora et al. 2024), including a documented case in the leaf miner moth *Cameraria ohridella* (Dalíková et al. 2017).

The degeneration of the sex-limited chromosome (W or Y) in the heterogametic sex leads to imbalances in gene copy number, which can disrupt the stoichiometry of protein complexes and reduce organismal fitness and lethality (Veitia 2005). To counteract these effects, many species have evolved sex-specific regulatory strategies (referred to as dosage compensation) that restore equilibrium in gene expression between sex chromosomes and autosomes, and consequently between the sexes (Lucchesi et al. 2005). These regulatory mechanisms may operate across the entire sex chromosome or target only specific, dosage-sensitive genes, resulting in either chromosome-wide or localized compensation respectively (Gu and Walters 2017). Different organisms have evolved diverse strategies that typically involve epigenetic regulation to achieve dosage compensation (DC) (Muyle et al. 2021)). In XX/XY systems, DC typically involves a chromosome-wide modulation of gene expression. For instance, in both placental mammals and *C. elegans*, DC is accomplished by repressing transcription in the homogametic (XX) sex (Lau and Csankovszki 2015). In contrast, *Drosophila* achieves DC by upregulating transcription from the single X chromosome in males to match the expression levels of the two X chromosomes in females (Samata and Akhtar 2018). This hyperactivation is driven by the deposition of activating histone modifications, notably H4K16ac, which promotes chromatin decompaction and facilitates increased transcription (Samata and Akhtar 2018).

ZW species were initially believed to exhibit only partial dosage compensation, but recent studies in Lepidoptera (moths and butterflies) and Artemia have revealed chromosome-wide compensation mechanisms (Bett et al. 2025; Gu et al. 2019). In *Artemia franciscana*, a well-differentiated region of the Z chromosome in females (ZW) is enriched with the active histone mark H4K16ac, enhancing transcription to balance the two Z copies in males (ZZ) (Bett et al. 2025; Zimmer et al. 2025). By contrast, the Lepidoptera DC mechanism works through dampening of expression of both Z chromosomes in males, reminiscent of the mechanism of nematodes (Rosin et al. 2022; Tomihara et al. 2022). In the monarch butterfly, gene expression along the Z chromosome shows a dichotomy of dosage compensation mechanisms: the ancestral Z is depleted of H4K16ac in males, in line with the nematode-like compensation of other lepidopterans, while the neo-Z is enriched with H4K16ac in females, resembling instead the mechanism of Drosophila (Gu et al. 2019). This dichotomy is surprising in light of various studies of Drosophila neo-sex chromosomes that show that the ancestral mechanism of dosage compensation is typically recruited to neo-X chromosomes through the acquisition of novel binding sites for the dosage compensation complex (Alekseyenko et al. 2013). Studies of additional neo-sex chromosomes are needed to shed light on whether there is a consistent difference between Lepidoptera and Drosophila, and what might be causing it.

In this study, we applied genomic approaches to investigate the evolutionary dynamics and regulatory architecture of dynamic multiple sex chromosomes in the horse-chestnut leaf miner, *Cameraria ohridella*. We sequenced, assembled, and annotated the genome of *C. ohridella* and utilizing previously published male and female genomic data, we assess the extent and structure of sex chromosome differentiation. To further explore the functional consequences of this differentiation, we analyze RNA-seq data from head and whole-body tissues, focusing on patterns of dosage compensation across distinct regions of the Z chromosome that were partitioned based on their evolutionary origins. We also examine the influence of sex-biased gene expression on dosage compensation dynamics. Finally, we investigate the chromatin landscape across these Z-linked regions using CUT&Tag profiling of key histone modifications; H4K16ac, H3K4me3, and H3K27me3 to probe their role in mediating dosage compensation.

## Results

### Complex Evolutionary Dynamics of Sex Chromosomes

In order to construct a high-quality genome for the female *Cameraria ohridella*, we generated and assembled 85.7 Gb of long-read PacBio HiFi data (∼214-fold coverage). The final contig genome spans approximately 453 Mb, comprising 447 contigs with an N50 of 2.5 Mb (Table S1). Scaffolding using consensus HiFi data resulted in an N50 length of ∼9.1 Mb. Assessment of genome completeness using BUSCO (*arthropoda_odb10*) revealed 98.4% completeness, including 92.2% single-copy orthologs (supplementary fig. 1), underscoring the high quality and completeness of the assembly. We focus here on the largest 34 scaffolds (referred to as superscaffolds 1-34), which contain 80% of the genome and 87% of BUSCO genes. Previous research has identified the presence of a neo-Z chromosome in *Cameraria ohridella*, formed by the fusion of the ancestral Z chromosome with autosomal one (chromosome 18 in *Bombyx mori*) (Fraïsse et al. 2017). To investigate this further, we computed coverage using short-read male and female DNA reads for 5,000 bp windows along the genome. Two superscaffolds, 3 and 30, had consistently low female:male coverage, consistent with being derived from the differentiated part of the Z chromosome (fig. 1b). These superscaffolds correspond to the ancestral Z and autosomal chromosome 18 in *Bombyx mori* as previously reported (Fraïsse et al. 2017). Both scaffolds map to the Z chromosome of *Euspilapteryx auroguttella* (fig. 1a and fig. 1c), a species within the same *Gracillariidae* family as *C. ohridella* (Boyes et al. 2024), showing that the fusion between the ancestral Z (AncZ) and the autosomal chromosome 18 (which we now refer to as NeoZ_1_) occurred prior to the divergence of the two species around 100 million years ago (Li et al. 2022).

**Fig 1:**
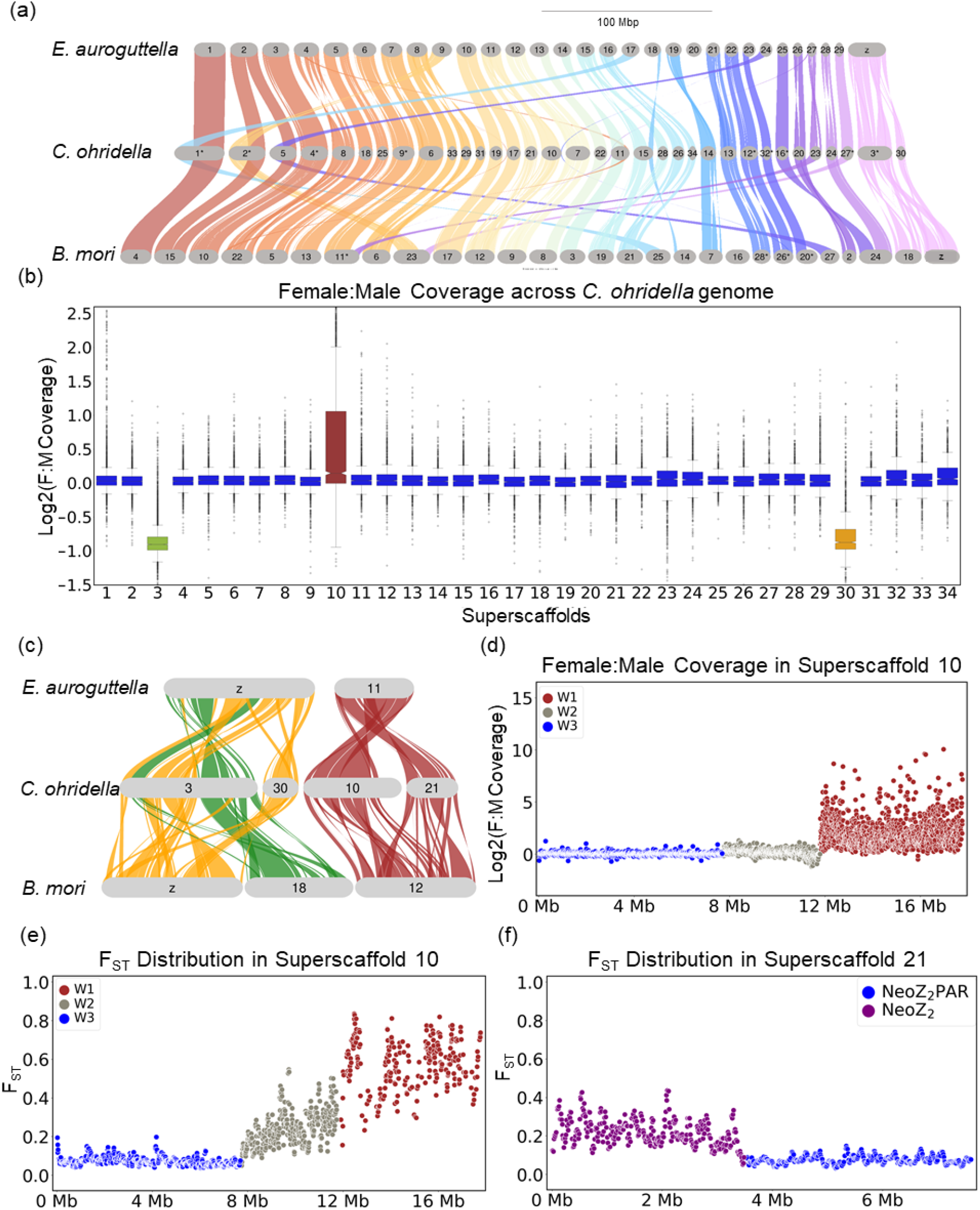
Identification and extent of differentiation of Z-linked and putative W-linked chromosomes. (a) Synteny across *Bombyx mori, Cameraria ohridella* and *Euspilapteryx auroguttella* (species within the Gracillariidae family) estimated using GENESPACE. (b) Female-to-male whole genome sequencing coverage over a 5kb window across the *Cameraria ohridella* genome assembly. (c) Synteny analysis of selected superscaffolds (sex chromosomes) of *Cameraria ohridella*, with distant related species *Bombyx mori* and closely related species *Euspilapteryx aurogutella* (same family as *C. ohridella*). (d) Female-to-male whole-genome sequencing coverage over a 5 kb window in superscaffold 10. (e) Female-to-male F_ST_ distribution over 10kb windows across superscaffold 10. (f) Female-to-male F_ST_ distribution over 10kb windows across superscaffold 21.

Superscaffold 10 had a strongly increased female:male coverage ratio compared to other chromosomes (fig. 1b), suggesting it may correspond to the W chromosome. We confirmed W-linkage using polymerase chain reaction (PCR) in five females and five males, with six out of 7 primer pairs consistently producing female-specific bands (supplementary fig. 2). A detailed examination (fig. 1d) showed various levels of female:male coverage along the scaffold, suggesting that parts of the W were acquired more recently than others. The presence of (female-specific) genetic variants on the W chromosome is often used to detect regions of ZW pairs that recently stopped recombining. We used genomic reads derived from pools of females and males to quantify genetic divergence (in the form of the fixation index F_ST_) between them. Integrating both coverage and F_ST_ analyses, we identified three distinct evolutionary strata within the W chromosome (figs. 1d and 1e). The first stratum (W1), spanning approximately 12Mb to 17Mb, is characterised by increased female coverage and elevated F_ST_ values, indicative of an older, highly differentiated W-linked region (figs. 1d and 1e). The second stratum (W2), extending from 8Mb to 12Mb, shows a moderate increase in female coverage combined with pronounced F_ST_, reflecting a more recent non-recombining region of the W chromosome. Finally, the segment from 0Mb to 8Mb exhibits similar coverage and F_ST_ values in both sexes (W3) (figs. 1d and 1e). The W chromosome shows several features typical of a degenerated sex chromosome. For instance, it has accumulated a remarkably high repeat content (79.42%), significantly exceeding the overall repeat content of the genome (56.84%) (supplementary fig. 3). This accumulation is accompanied by a low gene density (supplementary fig. 4) and reduced gene expression, with the median TPM of ∼0.4 being substantially lower than the autosomal median of ∼4 TPM. The W1 region maintains a slightly elevated GC content (38.32%) compared to the genome-wide average of 38.08% (supplementary fig. 4), consistent with reports of GC-rich, repeat-enriched W chromosomes, such as in *Dryas iulia*, where the W is approximately 15% more GC-rich than the rest of the genome and enriched for long repeats (LINEs and LTRs) (Lewis et al. 2021).

The origin of the W chromosomes of Lepidoptera is still under debate (Han et al. 2024). To investigate the origin of the W chromosome (superscaffold 10), we checked for sequence similarity between the genes on superscaffold 10 and the rest of the *C. ohridella* genome. Contrary to our expectation that most W genes should be homologous to Z or neo-Z-linked genes (superscaffolds 3 and 30), most genes from the W2 and W3 regions have homologs on superscaffold 21 instead (supplementary figs. 5 and 6). Furthermore, a synteny analysis showed that both superscaffold 10 (W) and superscaffold 21 are homologous to the *B. mori* chromosome 12 and *E. auroguttella* chromosome 11 (fig. 1c), suggesting that superscaffold 21 is a second neo-Z chromosome, which we refer to as NeoZ_2_. This was confirmed by an F_ST_ analysis of this scaffold, which revealed two strata: a region with elevated F_ST_ between males and females, consistent with a young non-recombining ZW region, and a distal region with uniformly low F_ST_ (fig. 1f, supplementary fig. 7).

Surprisingly, this analysis of genetic differentiation also identified three regions with elevated male:female F_ST_ values on the ancestral Z chromosome (AncZ) (figs. 2a and 2b), suggesting a recent duplication of parts of the Z to the W chromosome. Notably, a genome-wide homology analysis, in which all genes were compared against one another and filtered to retain their strongest matches, showed that these high-F_ST_ regions of the ancestral Z are homologous to each other (fig. 2d). We further mapped genes located in the high F_ST_ region on the AncZ to scaffolds that were not included in the primary superscaffolds (scaffolds 35 to 220) to search for putative W scaffolds with genes homologous to those in these genomic regions. Fourteen scaffolds were recovered, of which scaffolds 48, 50 and 51 had pronounced levels of female-to-male coverage and/or high male:female F_ST_ (fig. 2c, supplementary fig. 8), supporting the presence of a duplicated region of the Z on the W chromosome .

**Fig 2:**
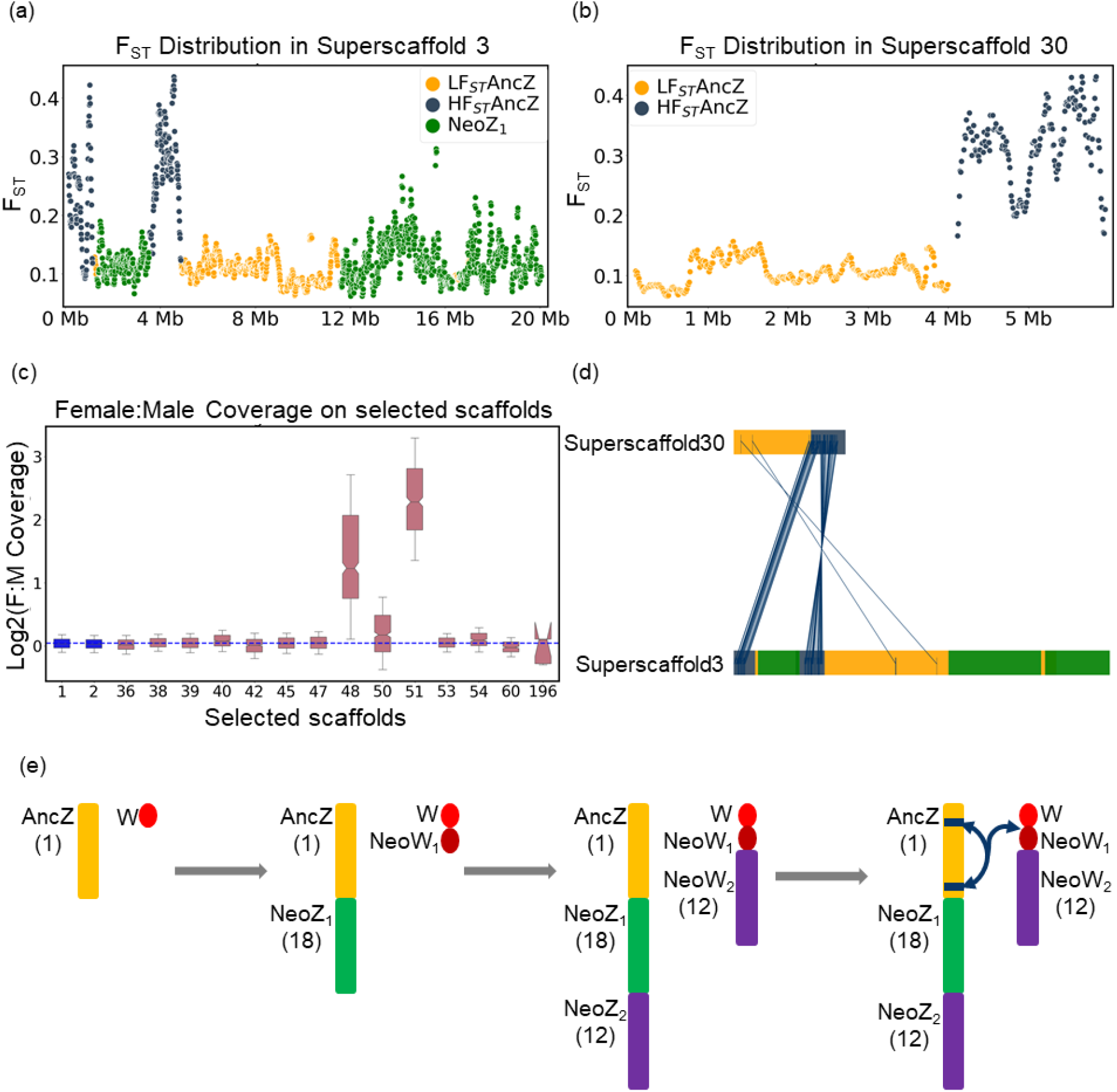
AncZ-W duplication and proposed model of evolutionary dynamics of sex chromosome evolution in *Cameraria ohridella*. (a). F_ST_ distribution across AncZ and NeoZ_1_ of the superscaffolds 3 (b). F_ST_ distribution across anc-Z of the superscaffolds 30 (c) Female-to-male coverage of putative W scaffolds selected based on gene homology with those in high-FST regions of AncZ. (d). High-F_ST_ regions of AncZ identified as homologous to one another through reciprocal best-hit analysis (e). This model proposes an evolutionary dynamic of sex chromosome evolution in *Cameraria ohridella* species with ZW system. Initially, there exists an ancestral Z (corresponding to chr1 in *B. mori*) and the W chromosome which is small and likely heterochromatic. Then, ancZ likely fused with an autosome (corresponding to chr18 in *B. mori*) resulting in a neo-sex chromosome (AncZ-NeoZ_1_) and its homologous pair (NeoW_1_) underwent rapid degeneration. The second fusion event occurred that involved W-NeoW_1_ with an autosomal chromosome (corresponding to chr12 in *B. mori*) creating a W-NeoW_1_-NeoW_2_. Concurrently, the Z chromosome acquired a homologous pair of chromosome 12, forming a chromosome AncZ-NeoZ_1_-NeoZ2. The final step is likely to involve ancestralZ-W duplication and structural rearrangements such as degeneration, translocation and gene duplications in the W. In summary, sex chromosome system consists of a large, multi-origin Z composed of ancestral and neo-sex chromosome regions (1, 18, and 12 corresponding chromosomes in *B. mori*), and a degenerate W chromosome containing ancestral W and portions of chromosome 12.

### Dosage Compensation in Sex Chromosomes with Dynamic Evolution

We examined gene expression in head and whole-body tissues to test whether dosage compensation occurred across the AncZ, NeoZ_1_, and NeoZ_2_ regions. Specifically, we tested, for each Z chromosome region: (1) whether male and female expression levels were equal (“dosage balance”), and (2) whether expression levels were broadly similar to those of autosomal genes (“complete dosage compensation”). In heads, genes on the AncZ showed significantly lower expression compared to autosomal genes in both sexes (*P*-value < 0.01, fig. 3a), in line with the mechanism of dosage compensation known in other Lepidoptera (which typically involves the down-regulation of male Z-linked genes (Kiuchi et al. 2014; Walters et al. 2015; Walters and Hardcastle 2011; Huylmans et al. 2017). Comparisons between males and females revealed a more complex pattern within the AncZ region: although overall expression was higher in females than in males (*P*-value=0.0001), this bias was largely driven by genes in high F_ST_ regions (*P*-value=1.9e-06, fig. 3c), suggesting that the female-bias in expression is due to the additional copies of these genes on the W chromosome. On the other hand, the genes in low F_ST_ regions exhibited balanced expression between the sexes (*P*-value=0.14, fig. 3c).

**Fig 3:**
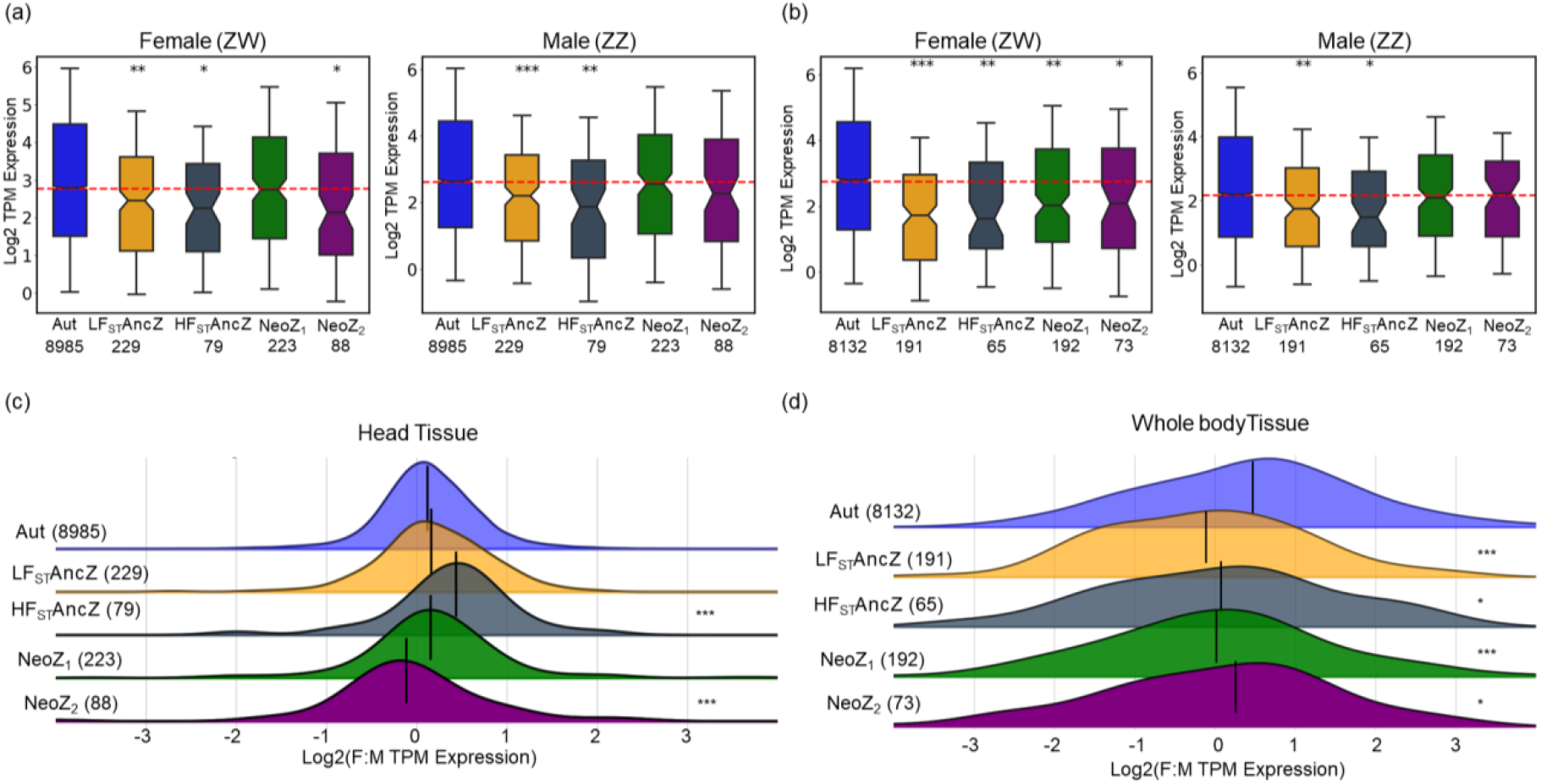
Patterns of dosage compensation between the Z-linked chromosomes and autosomes in heads and whole body tissues of both males and females. Distributions of Log_2_ transformed TPM expression values are given for two regions of ancestral Z (chr1 in *B. mori*), neo-Z_1_ (chr18 in *B. mori*), neo-Z_2_ (chr12 in *B. mori*) and autosomes for each sex and two tissues in Cameraria ohridella. The red dotted lines in box plots represent the overall expression median in each sex. The vertical black lines represent median values. Statistical significance of Wilcoxon signed-rank test applied to each data set between autosomes and each Z-linked region and indicated by stars (\**P*-value < 0.05, ** is *P*-value < 0.005, and *** is *P*-value < 0.0005).

NeoZ_1_ genes exhibited expression levels comparable to autosomes and showed no significant differences between males and females, suggesting complete dosage balance and compensation (*P*-value < 0.1, fig. 3a). The NeoZ_2_ showed reduced expression in females relative to males (*P*-value=0.0003, fig. 3c), suggesting that full dosage compensation has yet to evolve on this young sex chromosome. While patterns were qualitatively similar for the male whole bodies, female whole bodies showed a reduction in expression of genes in all Z-linked regions relative to autosomal genes (fig. 3c and d), in line with what has been observed in other Lepidoptera species when gonads are included in the analysis (Gu et al. 2017; Huylmans et al. 2017). The differential representation of sex-biased genes on sex chromosomes can bias the inference of dosage compensation from whole-body data, and their removal from such analyses has been recommended (Walters and Hardcastle 2011). When sex-biased genes (1790 autosomal genes, and 152 genes in the Z chromosome i.e. those with ≥2-fold expression differences between sexes and adjusted *P*-values < 0.05, supplementary fig. 9) were excluded in whole body tissues, the expression differences between males and females in NeoZ_2_ and high F_ST_ of AncZ were no longer statistically significant (supplementary fig. 10). However, the reduced expression of NeoZ_1_ and the low F_ST_ regions of AncZ in females persisted (supplementary fig 10). These findings highlight the role of tissue context and sex-biased gene content in shaping dosage regulation across sex chromosomes.

### Dual Epigenetic Dosage Compensation Mechanisms on the Z

In order to explore the chromatin landscape underlying dosage compensation along the Z chromosome in *Cameraria ohridella*, we examined the distribution of key histone modifications associated with transcriptional regulation. Specifically, we used CUT&TAG to measure the enrichment along the genome of H3K4me3, a mark of active promoters; and H4K16ac, an activating modification associated with transcriptional hyperactivation in the heterogametic sex in species such as *Drosophila, Artemia franciscana*, and the green anole. We also examined the repressive modification H3K27me3; however, unlike H3K4me3 and H4K16ac, which displayed distinct enrichment patterns associated with gene expression (supplementary figs. 11 and 12), H3K27me3 enrichment did not exhibit a clear correlation with expression levels (supplementary fig. 13) and therefore no further downstream analysis were conducted in this histone modification. Since the CUT&TAG analysis pipeline relied on uniquely mapped reads, high FST regions and NeoZ_2_ were not included in this analysis (as reads derived from these regions map to several locations in the genome).

To evaluate differences in chromatin activity between Z-linked and autosomal regions, we compared the distribution of active histone marks across genes and their ±500 bp flanking regions on the AncZ, NeoZ_1_, and autosomes in both sexes. In males, both Z-derived regions showed a significant reduction in H3K4me3 enrichment relative to autosomes (*P*-value = 0.02 for NeoZ_1_; *P*-value = 3.7e-10 for AncZ; figs. 4a and 4e), consistent with diminished transcription on the male Z. Average enrichment profiles across annotated genes (±1,000 bp around TSS and TES) revealed that this reduction was strongest near the TSS and gradually diminished toward the TES, where enrichment levels converged (fig. 4c). In contrast, females exhibited broadly comparable H3K4me3 levels across autosomal, AncZ, and NeoZ_1_ genes (*P*-value > 0.05, fig. 4a). However, H4K16ac exhibited a pronounced sex- and region-specific distribution. In females, this modification was markedly enriched on the NeoZ_1_, whereas the AncZ showed levels similar to those of autosomes (*P*-value = 7.1e-09 for NeoZ_1_, *P*-value = 0.12 for AncZ fig. 4b). In males, the pattern was reversed: the AncZ was significantly depleted for H4K16ac, while the NeoZ_1_ retained autosome-like levels (*P*-value = 0.71 for NeoZ_1_, *P*-value = 1.1e-16 for AncZ; fig. 4b) and this depletion was not only at TSS but also across the gene body length (fig. 4d). Although H4K16ac was significantly enriched on the NeoZ_1_ in females, genome-wide comparisons revealed no distinct region where NeoZ_1_ stood out relative to the rest of the genome or to males. This suggests that the observed NeoZ_1_ enrichment represents a subtle, rather than uniquely localized, chromatin difference (fig. 4f).

**Fig 4:**
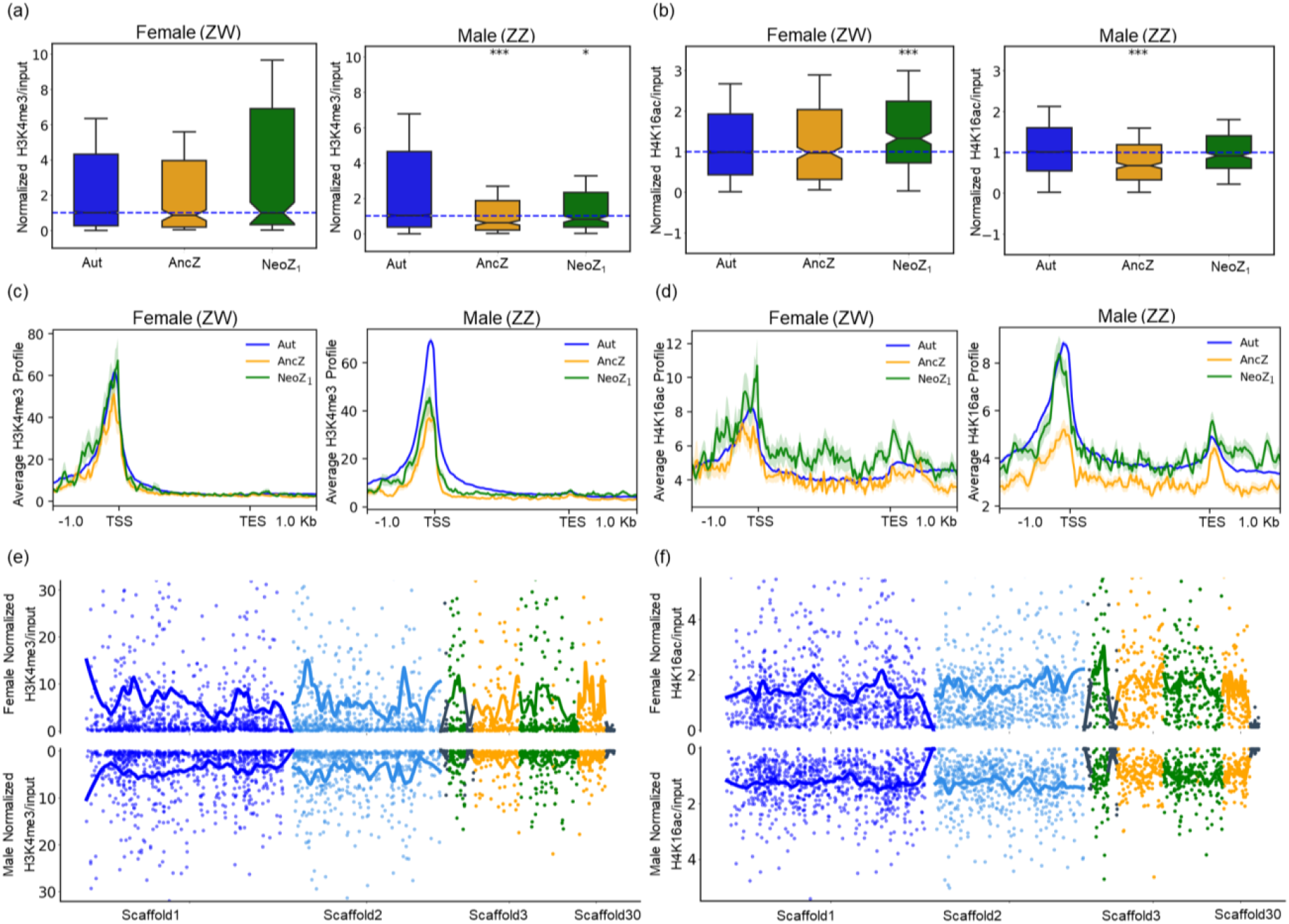
Chromatin landscape between the Z chromosome and autosome in head tissues of males and females of *Cameraria ohridella*. (a–b) Average enrichment profiles of H3K4me3 and H4K16ac across gene bodies and ±1 kb flanking regions. c–d) Normalized H3K4me3 and H4K16ac enrichment across genes located on the two Z-linked regions (AncZ and NeoZ_1_) compared with representative autosomes (scaffolds 1 and 2). Vertical black lines indicate median values. Statistical significance was assessed using Wilcoxon signed-rank tests (*P* < 0.05, P < 0.005, *P < 0.0005). e–f) Genome-wide distribution of H3K4me3 and H4K16ac across representative autosomal superscaffolds (1 and 2) and Z-linked regions (AncZ and NeoZ_1_).

## Discussion

Here, we uncover the evolutionary trajectory of a complex multi-sex chromosome system in *Cameraria ohridella*. This configuration is reminiscent of the multiple sex chromosome systems described in *Danaus chrysippus* (Martin et al., 2020; Smith et al., 2016) and *Ivela similis* (Mora et al., 2024), and generally in line with the many neo-Z chromosomes that have been described in Lepidoptera (Wright et al. 2024; Yoshido et al. 2020; Nguyen et al. 2013). It is unclear why such sex chromosome autosome fusions arise, but a role of sexually antagonistic selection has often been invoked: if a fusion (partly) links a female-beneficial male-deleterious allele and the female-determining locus, it will be favored by selection (Anderson et al. 2020). One key prediction of this hypothesis is that fusions that involve sex chromosomes should be more common than expected given the number of autosome/autosome fusions. Interestingly, while this seems to be the case in some clades, the opposite is found in others (such as Drosophila) (Anderson et al. 2020). Lepidoptera seem like a promising clade to investigate this given the large amounts of genomes that have been produced (Wright et al. 2024), and their variety of sex chromosome configurations. How sex chromosomes are regulated may also contribute to this diversity, as the adoption of ancient dosage compensation mechanisms by new sex chromosomes may mitigate the fitness effects of W/Y chromosome degeneration. However, this effect is hard to test as in most species how neo sex chromosomes are regulated is not known. Studies such as ours, where both neo sex chromosomes and their regulatory mechanisms are described, are a key step towards overcoming this limitation.

We observed an enrichment of H4K16ac on the NeoZ_1_ in females, whereas this mark is depleted on the AncZ in males, indicating the independent evolution of distinct dosage compensation mechanisms across the two Z segments. This mirrors the pattern seen in *D. plexippus*, although it involves the fusion of a different autosome (chromosome 16) with the ancestral Z (supplementary fig. 14). This is at odds with the well-studied situation of Drosophila neo-sex chromosomes, which largely appear to progressively co-opt the ancestral mechanism of dosage compensation for their regulation (Alekseyenko et al. 2013; Marín et al. 1996). Why should some mechanisms of compensation be co-opted but not others? One possible explanation for this difference is that when the ancestral mechanism of compensation involves the reduction of expression in the homogametic sex, its co-option for a newly evolved sex chromosome is not favored by selection, as it would spread the X/Z:autosome expression imbalance to both sexes (Toups and Vicoso 2025).

To illustrate this idea, we ran simple SLIM simulations of the evolution of a short regulatory sequence (200bps) controlling the expression of an X-linked gene having recently lost its Y counterpart (i.e. males still had reduced expression, and this caused a reduction in fitness). Mutations that modify expression (cis-eQTLs) could either have the same effect on males and females, or act specifically in one sex. In the absence of an ancestral chromosome-wide mechanism of compensation, male expression increased slowly, with only a minority of simulations reaching 90% of the optimum in 20000 generations. Allowing for the accumulation of dosage compensation complex (DCC) binding sites that reduced expression in the homogametic sex (“DC in Hom.” in fig. 5) did not substantially modify these dynamics, and such binding sites were never fixed, supporting the idea that ancestral down-regulation in the homogametic sex can be difficult to co-opt. On the other hand, binding sites for a DCC that increased expression in the heterogametic sex (“DC in Het.”) were fixed in 100% of the simulated regulatory sequences, and led to the fast equalization of expression between the sexes, in line with Drosophila neo-X chromosome evolution.

**Fig 5:**
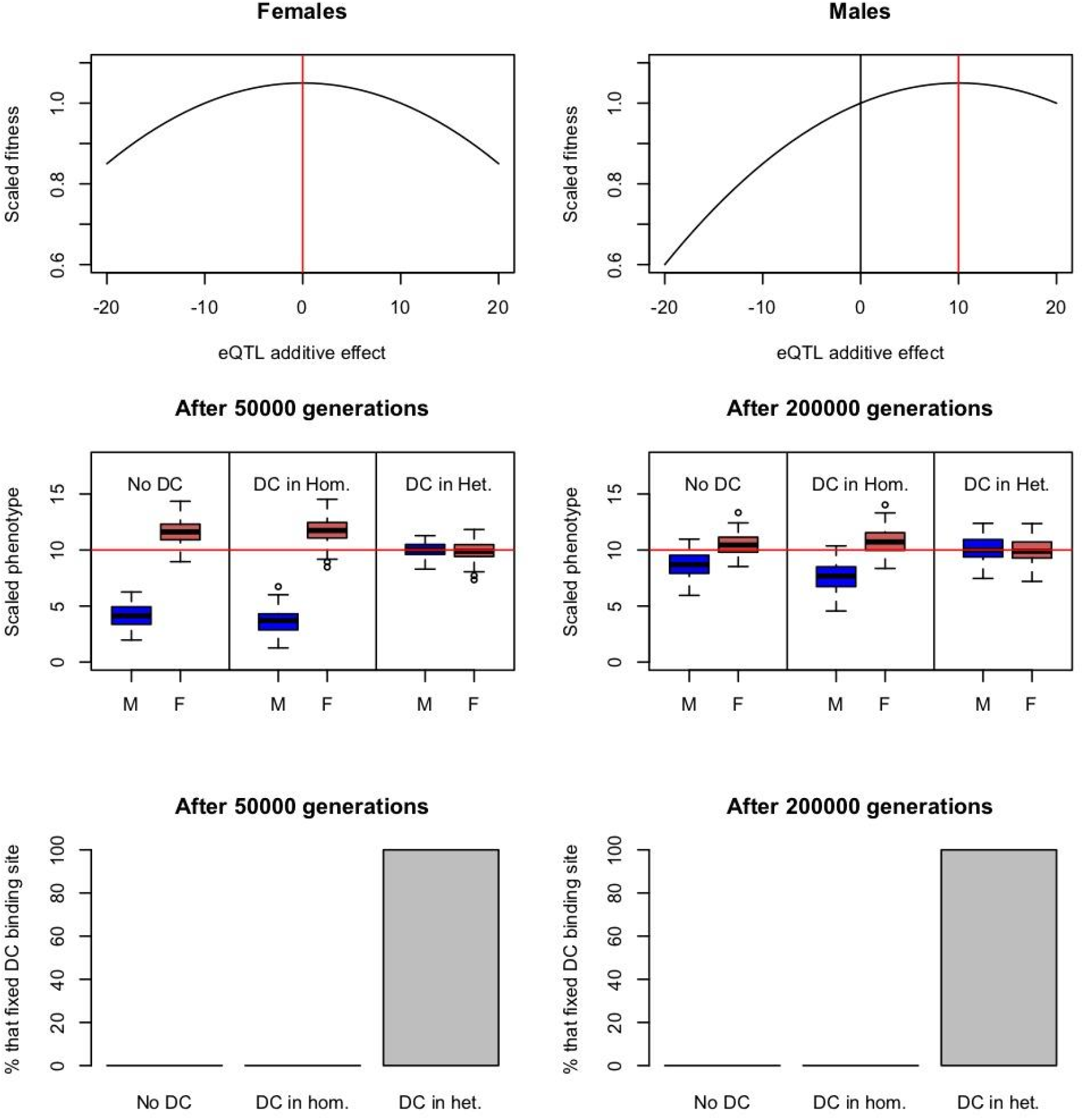
Simulations of the evolution of expression of a hemizygous neo-X-linked gene under different ancestral modes of dosage compensation. Upper panels: scaled male and female fitnesses as a function of the additive effect of all eQTLs found in each individual. The red line indicates the optimum, and the black line the starting point for males (XX females already start at the optimum). Middle panels: Gene expression evolution after 50000 and 20000 generations. The y-axis (Scaled phenotype) represents mean expression across the population, with the red line representing the optimum. M stands for Male and F for Female. Bottom panels: The proportion of simulations of each type where a binding site for dosage compensation was fixed. Each type of simulation was run 100 times.

While meant to be illustrative rather than realistic, these simulations also highlighted some situations that could favor the cooption of an ancestral mechanism that reduced homogametic sex expression (supplementary fig. 15). When no sex-specific eQTLs were allowed, a minority of simulations (4%) fixed DCC binding sites. If DCC binding site fixation only caused a partial down-regulation (-4 instead of -10), >40% of regulatory sequences acquired them. When both of these conditions were true, all simulations ended with fixed DCC binding sites. This suggests that the regulatory architecture of compensation will influence whether it gets co-opted by new sex chromosomes, with more subtle mechanisms being favored when the ancestral mechanism works through downregulation of the homogametic sex. Within a single genome, genes whose regulatory architecture is tightly coupled between the sexes may also be more likely to co-opt such ancestral mechanisms. Over 30 Z-autosome fusions have now been detected in Lepidoptera (Wright et al. 2024) with many more likely to follow with the generation of more chromosome-level genome assemblies. An in-depth investigation of their transcriptional and chromatin profiles, as well as that of neo-sex chromosomes in other clades with different ancestral modes of compensation, will allow for the direct testing of these hypotheses.

## Materials and Methods

### DNA Extraction and Sequencing

We collected 30 *Cameraria ohridella* females from European horse-chestnut trees in Vienna, Austria for high molecular weight DNA extraction using the Qiagen Genomic-Tip 20/G Kit. The DNA was sequenced on a PacBio Sequel II SMRT cell at the Vienna Biocenter sequencing facility.

### Genome Assembly and its Quality Assessment

The consensus sequences from the reads were assembled using Hifiasm with default parameters v0.19.9 (Cheng et al. 2021). The primary assembly was then processed to remove duplicates with consensus reads, using purged_dups over four iterations (Guan et al. 2020). To scaffold the purged assembly, we employed LongStitch (ntlink + arcs) (Coombe et al. 2021). We then assessed the completeness of the genome assembly using BUSCO v5.2.2 with the arthropoda_odb10 database (n = 1,013 single-copy orthologs) (Manni et al. 2021). To further evaluate genome completeness based on k-mer representation, we used Merqury v1.4, which compares the assembled genome against k-mers derived from Illumina short reads. K-mer databases were generated using Meryl v1.4, and completeness evaluation was performed in Merqury using default settings, providing insights into the representation of the nuclear genome.

### Coverage and F_ST_ Analysis

Male and female Illumina short-read genomic data, as described by Fraïsse et. al 2017 (Fraïsse et al. 2017), were aligned to the reference genome using Bowtie2 (v2.4.5) with the --end-to-end and --sensitive parameters (Langmead and Salzberg 2012). To retain only uniquely mapped reads, SAM files were filtered to exclude alignments containing the “XS” tag, which indicates secondary alignments, using grep -vw “XS”. Genome-wide coverage was then computed in non-overlapping 5 kb windows using the soap.coverage tool. To investigate population differentiation between sexes, we used aligned sequencing reads from two pooled DNA libraries; each representing 10 male and 10 female individuals. Alignments with a mapping quality (MAPQ) score below 20 were excluded and the remaining alignments were subsequently sorted using the view and sort functions of samtools v1.18 (Li et al. 2009). We then generated a pileup file of the filtered male and female alignments using the mpileup function in samtools. To quantify genetic differentiation, we employed the Popoolation2 pipeline (Kofler et al. 2011). A synchronized (sync) file was produced using the sync module of Grenedalf v0.2.0 (Czech et al. 2024) and used as input for Popoolation2’s fst-sliding.pl script. We estimated F_ST_ values between male and female pools across the genome using non-overlapping sliding windows of 10 kb. The analysis was conducted with the following parameters: --suppress-noninformative --min-count 5 --min-coverage 20 --max-coverage 2000 -- min-covered-fraction 0.0 --window-size 10000 --step-size 10000 --pool-size 20

### Repeat Annotation and Gene Prediction

A *de novo* consensus repeat library was generated using RepeatModeler v2.0.4 (Flynn et al. 2020), and repetitive elements were annotated across the genome with RepeatMasker v4.1.6 (Tarailo-Graovac and Chen 2009). To quantify repeat content, the genome was divided into non-overlapping 10,000 bp windows, and a custom Python script was used to calculate the proportion of repetitive elements from the RepeatMasker output. GC content was estimated for the same window size using the ‘-pattern GC’ option in bedtools nuc. Gene prediction was performed on the soft-masked genome using BRAKER v3.0.8 (Hoff et al. 2019), incorporating RNA-seq data from head and whole-body tissues, along with protein sequences from OrthoDB v12 (Arthropoda). The resulting gene models were refined using AGAT to remove transcript isoforms, incomplete genes lacking start and/or stop codons, and open reading frames shorter than 100 bp. Protein and coding sequences were extracted with the getAnnoFastaFromJoingenes.py script. The completeness of the annotated gene set was assessed using BUSCO v5.2.2.

### RNA-seq Processing and Differential Expression Analysis

*Cameraria ohridella* specimens that were collected from European horse-chestnut trees in Vienna, Austria, between May and September 2024, were sexed, identified and stored at -80 °C until their subsequent processing. RNA was extracted from adult male and female specimens, with each biological replicate comprising either pooled heads (five individuals) or a single whole-body sample. RNA-seq libraries were prepared and subjected to high-throughput sequencing. Raw reads were pseudoaligned to the annotated coding sequences (CDSs) of *C. ohridella* using Kallisto with default parameters (Bray et al. 2016) to estimate transcript abundances, reported as transcripts per million (TPM) and estimated counts. Differential gene expression (DGE) analysis was performed in R using the DESeq2 package (v4.3.2) (Love et al. 2014). A likelihood ratio test (LRT) framework was employed to assess sex-biased transcriptional differences within each tissue. Genes were considered significantly differentially expressed if they exhibited a log2 fold change >1 (i.e., >2-fold change) and a Benjamini-Hochberg-adjusted false discovery rate (FDR) <0.05. Differentially expressed genes were further categorized as male-biased or female-biased based on the direction of expression change.

### Identification of W-linked Candidates and PCR Validation

We implemented a k-mer-based pipeline using the BBMap tool (Bushnell 2014b), following the approach described by (Elkrewi et al. 2021), to identify female-specific transcripts. It involves first the generation of short k-mer sequences from both male and female DNA and RNA reads. Then k-mers with matches in male DNA and RNA were filtered out, and RNA reads that matched female-specific k-mers were extracted and were subsequently assembled. The resulting transcripts were then screened for genomic homology using pblat (Wang and Kong 2019). Regions within superscaffold 10 (putative W chromosome) that showed homology to these candidate W-linked transcripts were identified. Based on these loci, primers were designed for seven candidate sequences using Primer3 Plus (Untergasser et al. 2012). The links for primer sequences and the corresponding positions of their target regions within superscaffold 10 are provided in the supplementary data.

### CUT&Tag Library Preparation and Processing

For each replicate, head tissues from four field-collected adult individuals were pooled and permeabilized for 30 minutes in an ice-cold solution containing 10 mg collagenase in 1 ml Enzyme Dissociation Buffer. CUT&Tag libraries were then prepared using the Active Motif Kit, following the protocol described (Bett et al. 2025). We also checked for the quality for the paired end reads using fastqc (Andrews 2010). Overrepresented sequences were filtered out using the bbduk.sh package of bbmap tool (Bushnell 2014a)with the options “*ref = overrepresented*.*fasta k=25 hdist=1*.” Sequencing adapters were then trimmed with Trim Galore under parameters *“--paired - -nextera -q 30 --length 20*”. The trimmed paired-end reads were aligned to *C. ohridella* reference genome using bowtie2 v2.4.5 (Langmead and Salzberg 2012)with these options: *“--local --very-sensitive --no-mixed --no-discordant --phred33 -I 10 -X 700*.” The resulting alignments were filtered to remove secondary alignments with the options “grep -v ‘XS:i:’”. To generate average plots of chromatin signals around genes for each replicate, we used deepTools/bamCompare (Ramírez et al. 2016) to calculate normalized coverage in 20-bp bins “*bamCompare -b1 sample*.*bam -b2 input*.*bam --normalizeUsing RPKM --scaleFactorsMethod None --binSize 20 -- operation ratio --minMappingQuality 30 --minFragmentLength 50*”. The resulting files were processed with deepTools/computeMatrix, and the matrices were visualized using the plotProfile function.

### Simulations

Simulations were adapted from section “13.2 A simple model of variable QTL effect sizes” (p. 282 of the SLIM manual, version 5.1, Haller et al. 2026). We simulated a 200bp regulatory sequence controlling the expression of a gene on a male-haploid X, right after it lost its Y homolog (i.e. expression is lower in males than females). Expression values were under stabilizing selection in females, i.e. these were at the optimum when the cumulative effect of eQTLS in the regulatory sequence was 0. Males were below the optimum, which they reached when the cumulative effect of eQTLs was +10.

The specific fitness functions that we used are:

female: 1.05 - (x - 10.0+10)^2^ * 0.0005

male: 1.05 - (x - 10.0)^2^ * 0.0005

Where x is the cumulative additive effect of eQTLs present in each individual. The population size was set to 1000 individuals, and the mutation rate to 1e-06 (an unrealistically high value to allow for enough mutations to appear given the small number of individuals). A recombination of 1e-07 was used.

In the simplest simulation mode (mode 1, no DC), only three kinds of eQTLs exist: 1. eQTLs acting equally on both sexes, whose effect is sampled from a normal distribution centered around 0 with standard deviation of 0.5. 2. eQTLs acting specifically in females. 3. eQTLs acting only in males.The effect of sex-specific eQTLs is also sampled from the same normal distribution.

In the second mode (“DC in Het.”), dosage compensation happens in the heterogametic sex (like in *Drosophila*). In addition to the eQTLs described above, new DC binding sites can accumulate in the regulatory region and immediately produce a +10 effect only in the heterogametic sex. In the third mode (“DC in Hom.”), DC happens in the homogametic sex (like in nematodes). In addition to the eQTLs described above, new DC binding sites can accumulate in the regulatory region and immediately produce a -10 effect only in the homogametic sex (but with a dominance coefficient of 0.5, such that a single copy will cause a decrease of -5). Finally, we ran mode 3 with two modifications: DC binding sites cause only a partial reduction in female fitness (-4 instead of -10, PARTIAL), or without sex-specific eQTLs (NOSEX). We ran each simulation mode 100 times for 20000 generations, and tracked both the final mean phenotypic values of males and females, and the proportion of genes that fixed a binding site.

Code for these simulations is available at: (https://git.ista.ac.at/bvicoso/bett26/-/blob/main/slim_simulations.md).

## Supporting information

https://seafile.ist.ac.at/f/e287ebd703034521acc0/

