## Supplementary material for "Out with the Old: Contrasting Histone Marks are Associated with Dosage Compensation on the Ancient and New Z of a Moth with Complex Sex Chromosomes": https://seafile.ist.ac.at/f/e287ebd703034521acc0/

### **Supplementary Information**

| Statistic | Contig-level | Scaffold-level |
| --- | --- | --- |
| N50 length (bp) | 2569677 | 9121241 |
| N80 length (bp) | 1142567 | 4614841 |
| Longest contig/scaffold length (bp) | 9193191 | 30143683 |
| Number of contigs/ scaffolds | 447 | 220 |
| Total size (bp) | 453825874 | 453869025 |

Table S1: Genome assembly statistics of female *Cameraria ohridella*

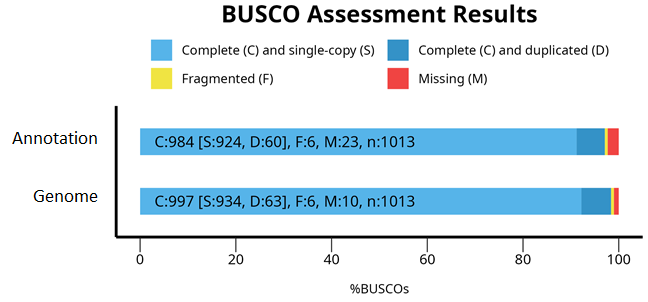

**Supplementary Fig. 1**: BUSCO assessment statistic results of genome assembly and annotation of female *C. ohridella*

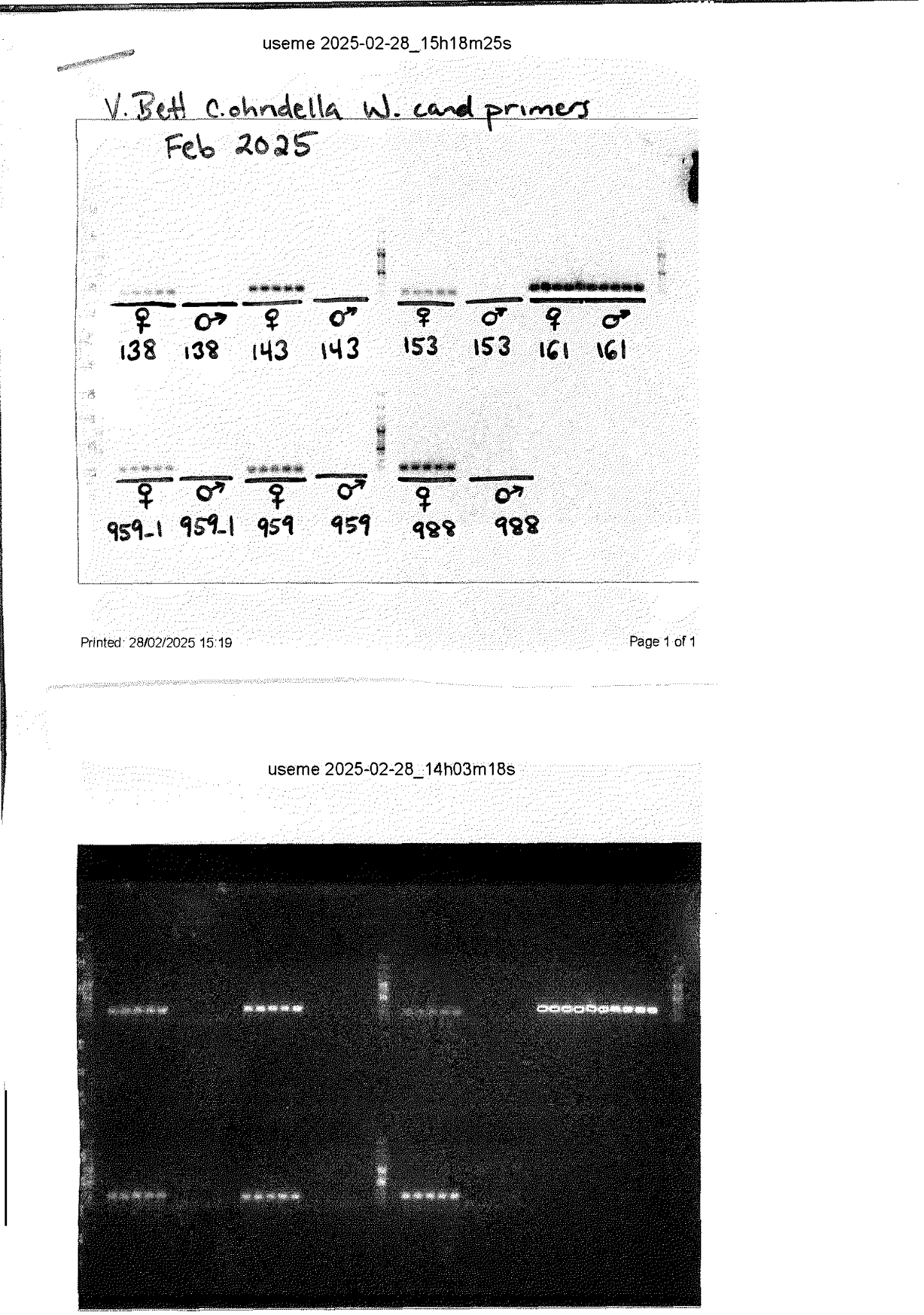

**Supplementary Fig. 2**: PCR for selected putative W-linked sequences (seven specific fragment sequences) conducted in five females and five males.

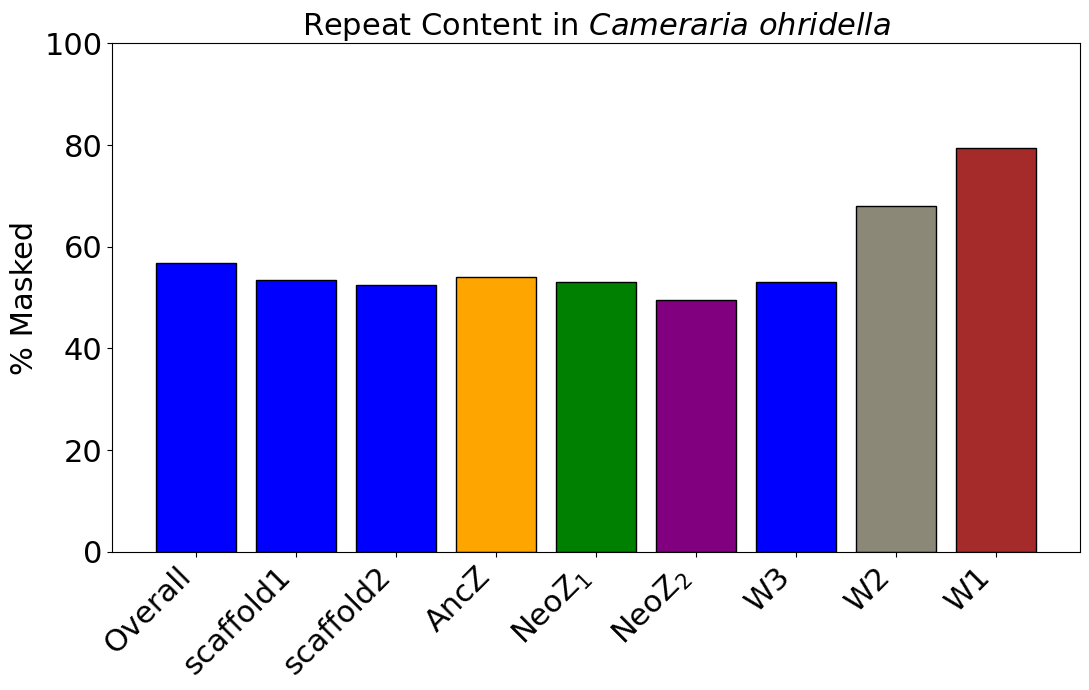

**Supplementary Fig. 3**: Repeat-masked regions across the genome, selected scaffolds, and sex-chromosome segments of *Cameraria ohridella*

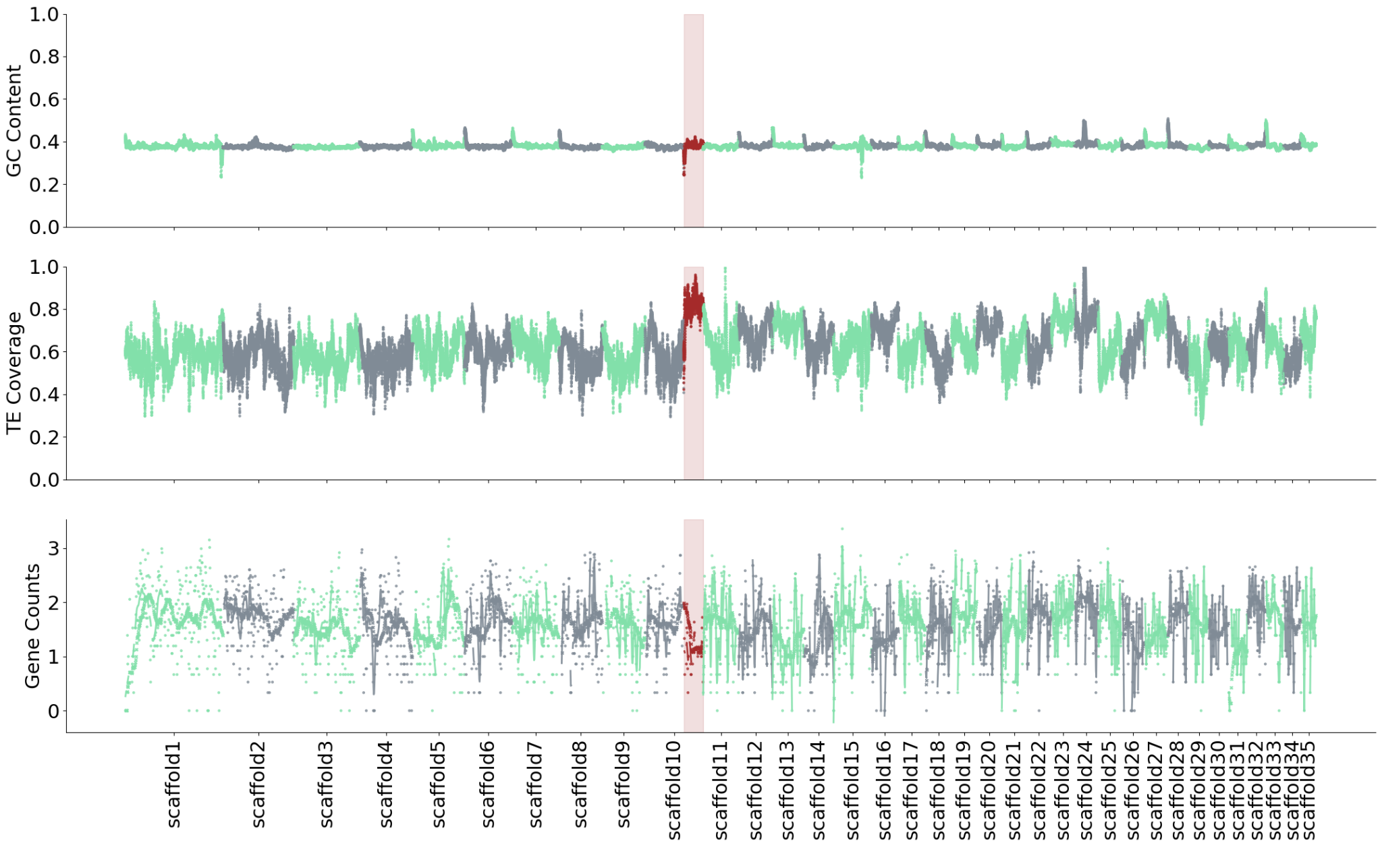

**Supplementary Fig. 4**: GC content, gene density and repeat content distribution across the genome of *Cameraria ohridella*

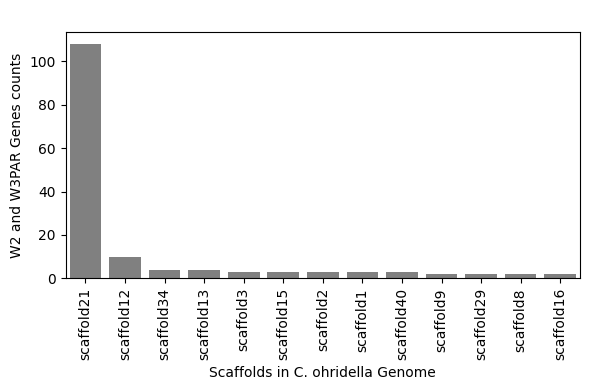

**Supplementary Fig. 5**: Distribution of genes on superscaffold 10 (W2 and W3, where repeat content is comparable to the overall genome) across *C. ohridella*.

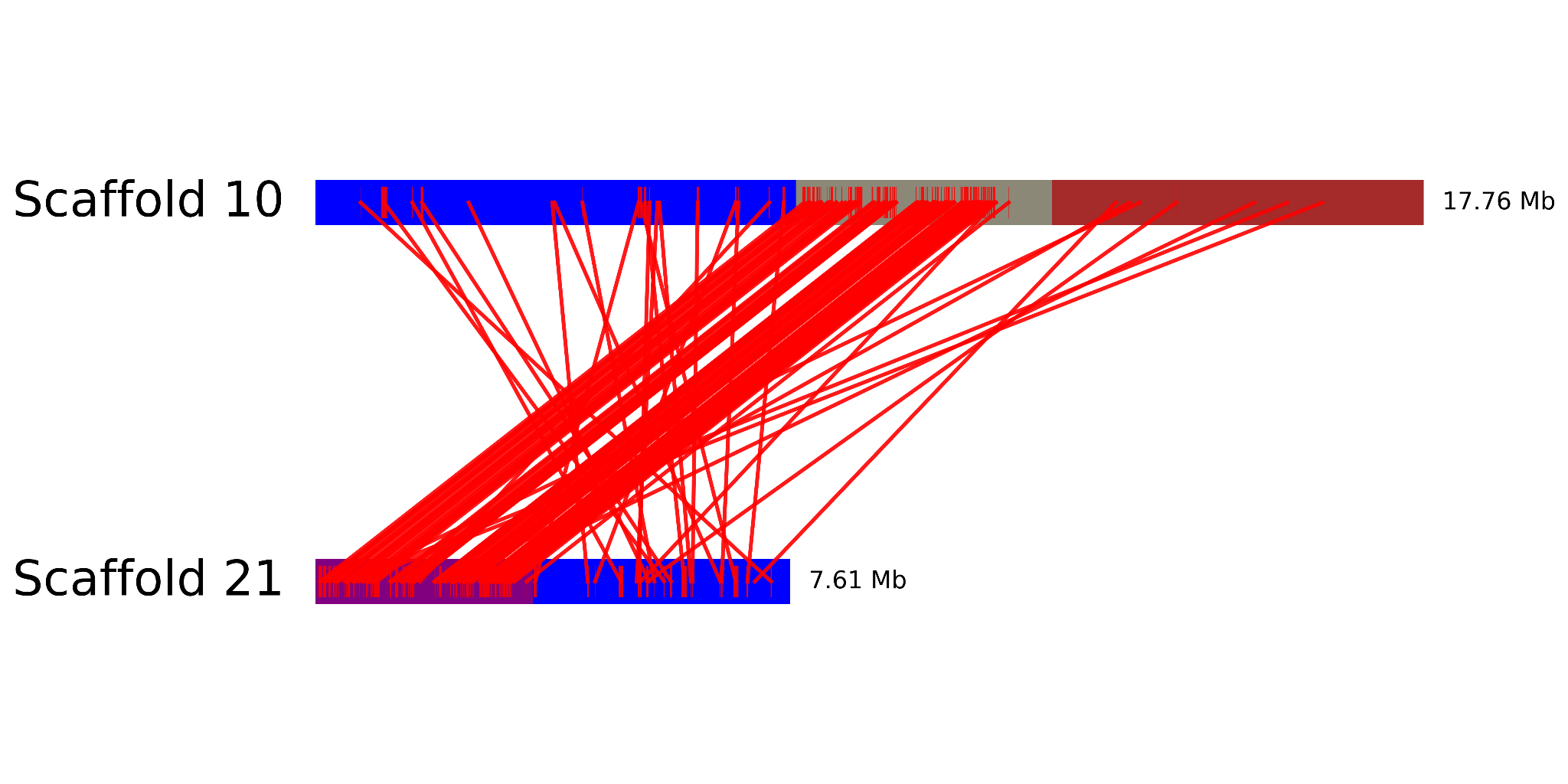

**Supplementary Fig. 6**: Synteny of superscaffold 10 and superscaffold 21

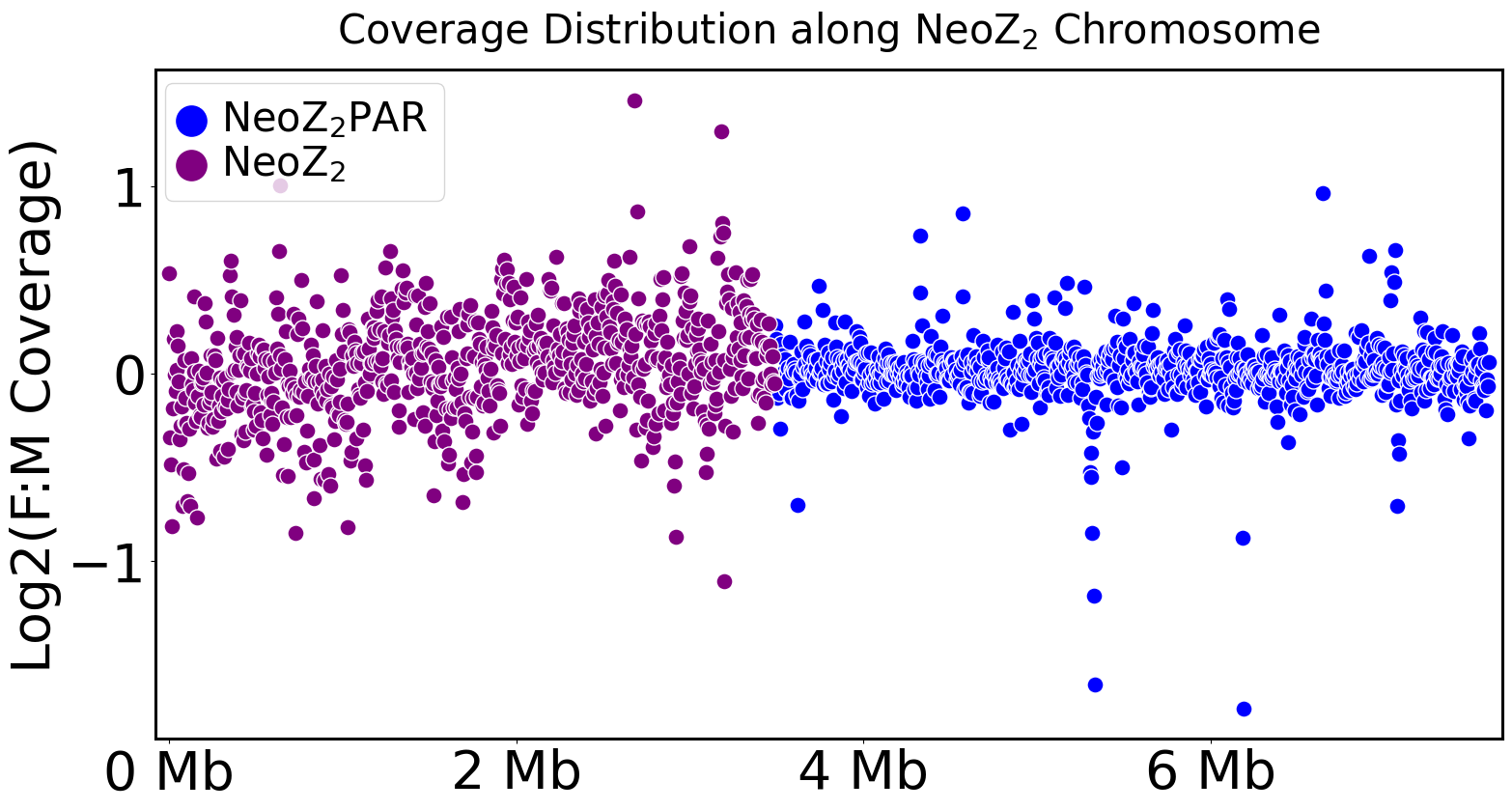

**Supplementary Fig. 7**: Female to male coverage between autosome to NeoZ_2_ in *Cameraria ohridella*

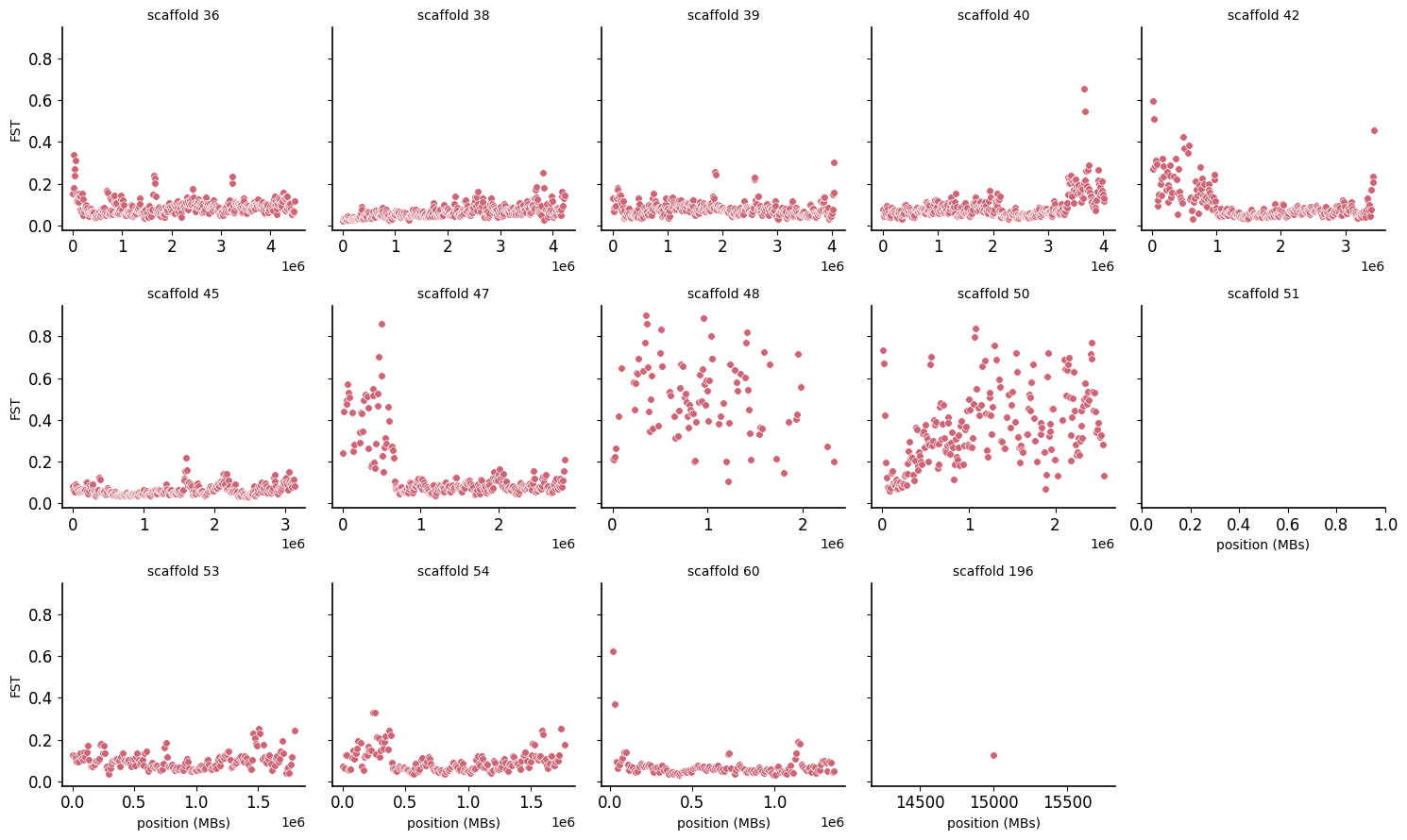

**Supplementary Fig. 8**: F_ST_ distribution across scaffolds with sequences homologous to HighF_ST_ regions of the ancestralZ

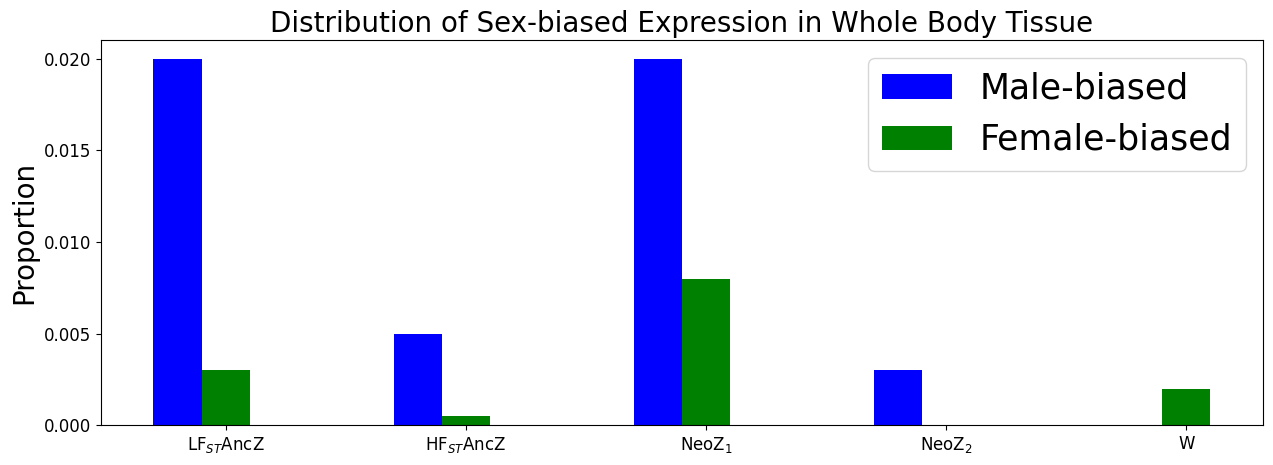

**Supplementary Fig. 9**: Overall sex-biased distribution in the Z chromosome (ancestralZ (LF_ST_ and HF_ST_ regions), NeoZ_1_, NeoZ_2_ and also in the W1-specific region of superscaffold 10.

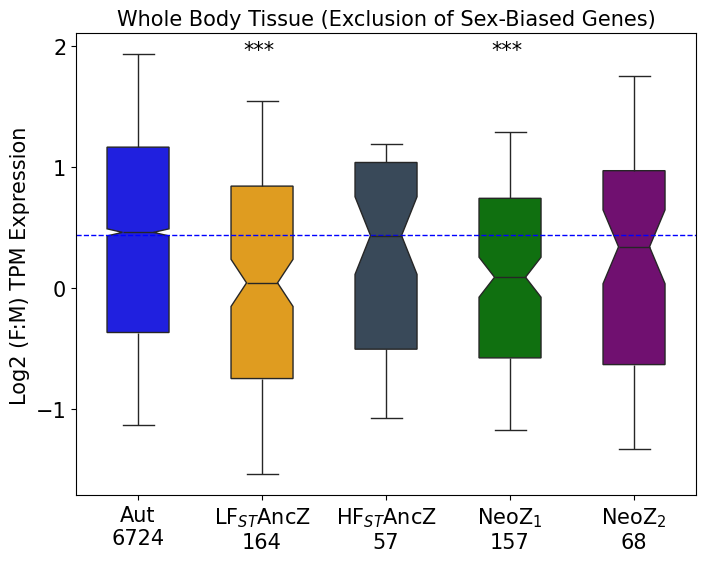

**Supplementary Fig. 10**: Patterns of dosage balance between the Z chromosome and autosome in whole body tissue following exclusion of sex-biased expression.

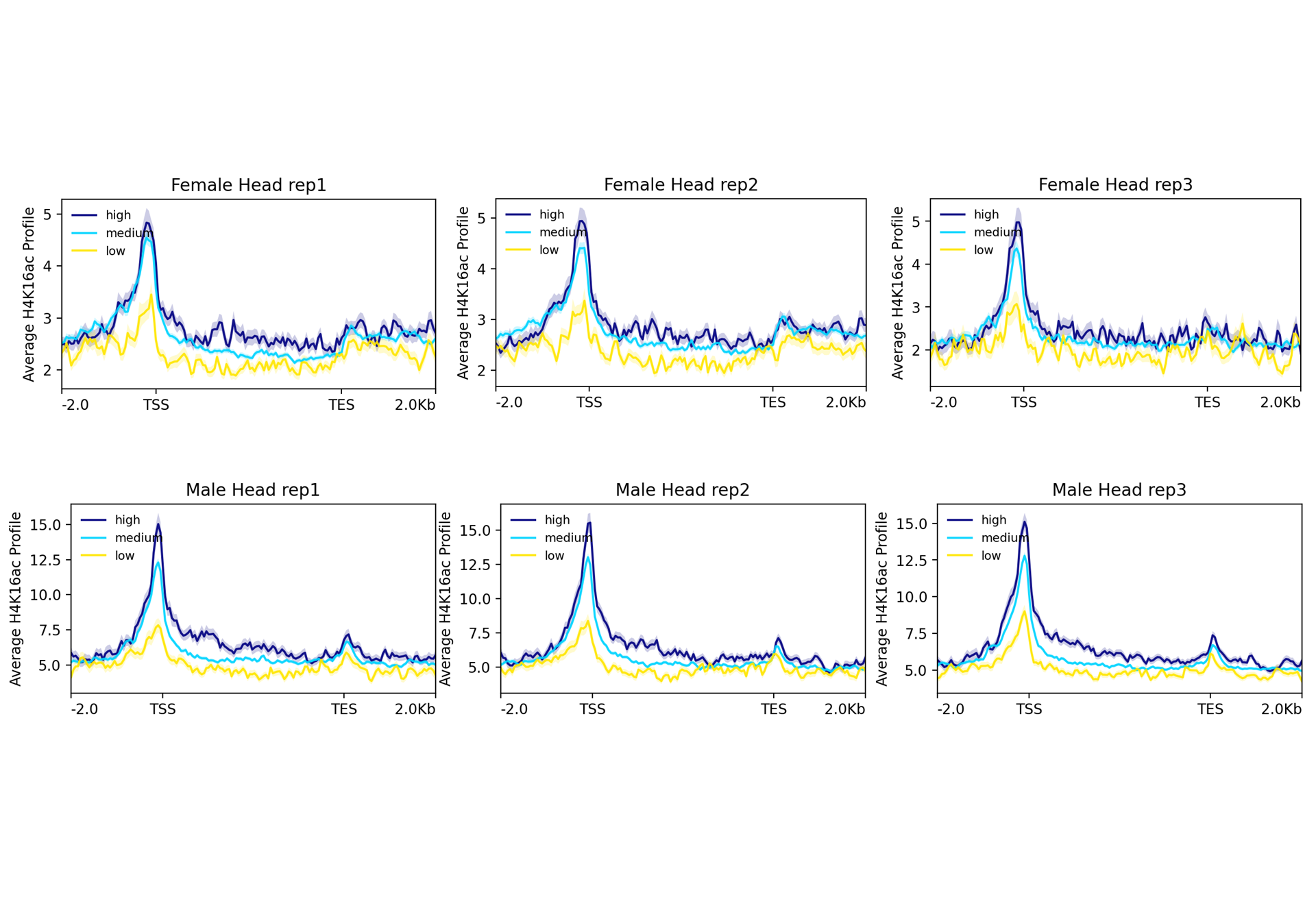

**Supplementary Fig. 11**: Correlation of enrichment of H4K16ac with expression levels

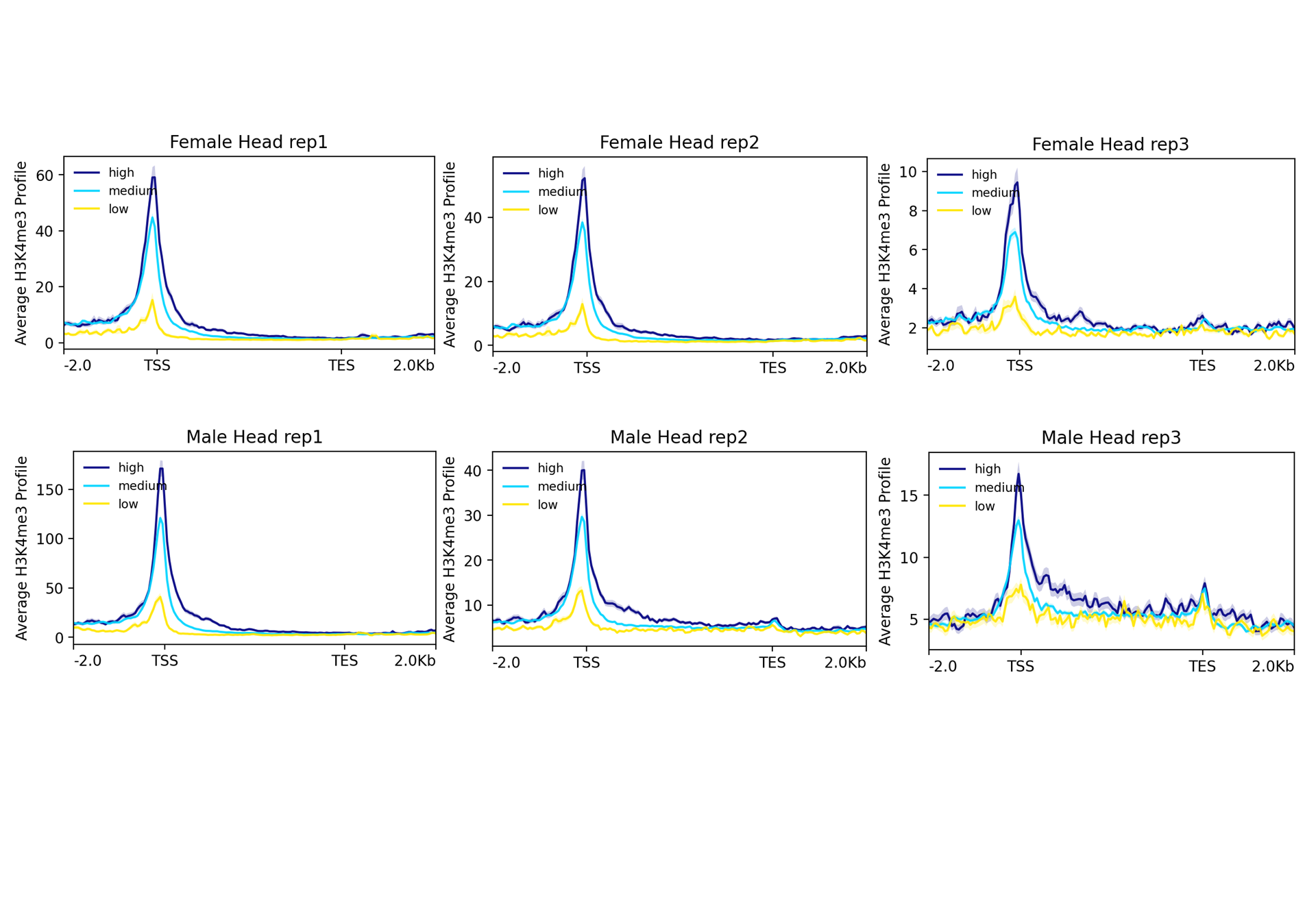

**Supplementary Fig. 12**: Correlation of enrichment of H3K4me3 with expression levels

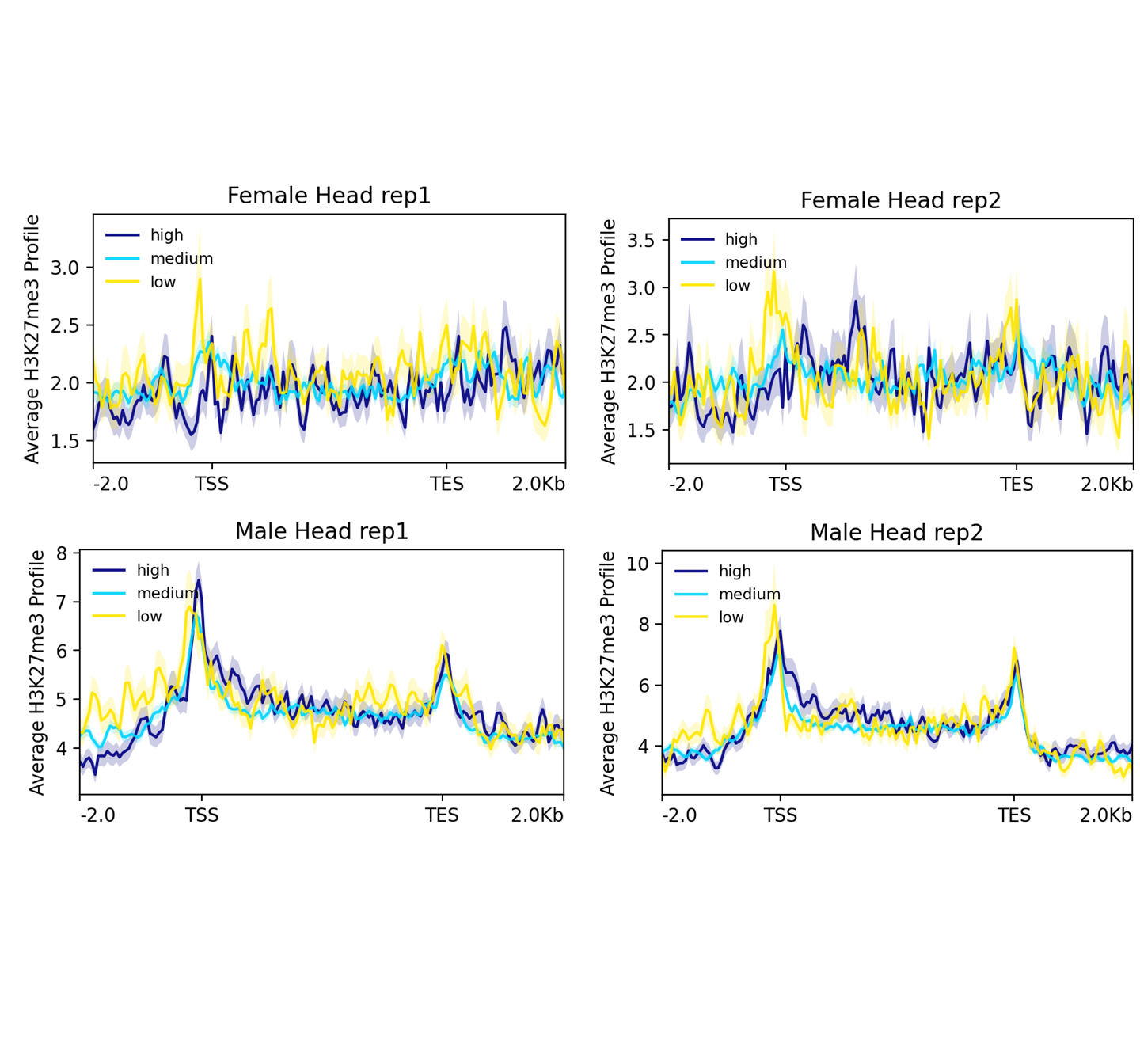

**Supplementary Fig. 13**: Correlation of enrichment of H3K27me3 and expression levels
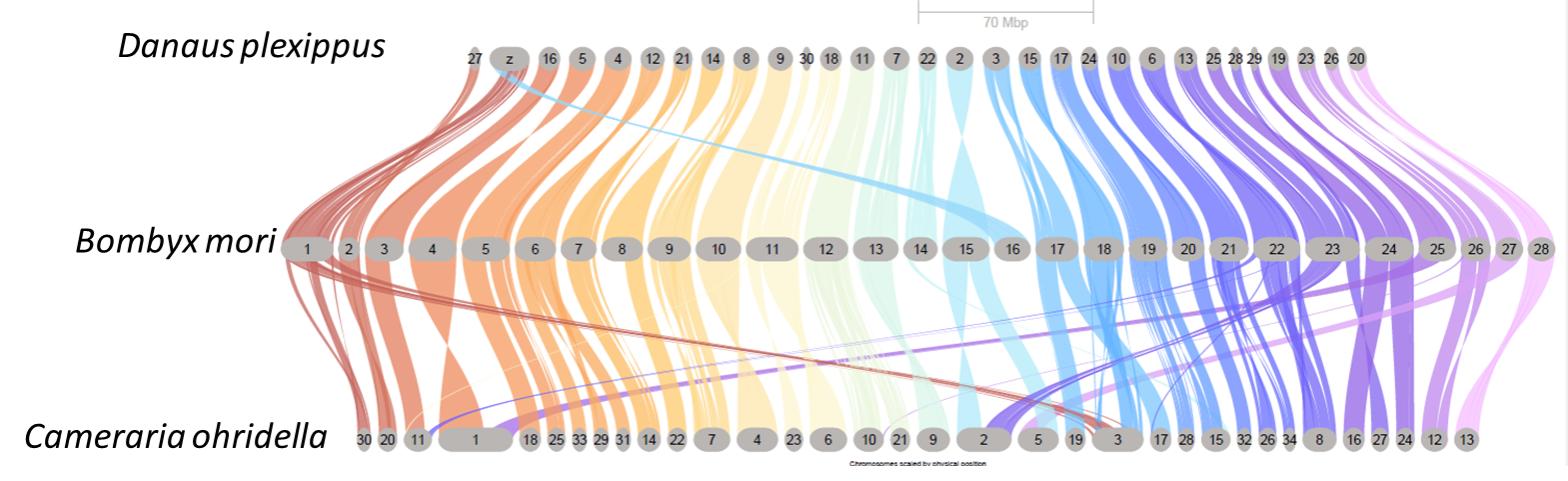

**Supplementary Fig. 14**: Synteny among *Danaus plexippus*, *Bombyx mori* and *Cameraria ohridella*

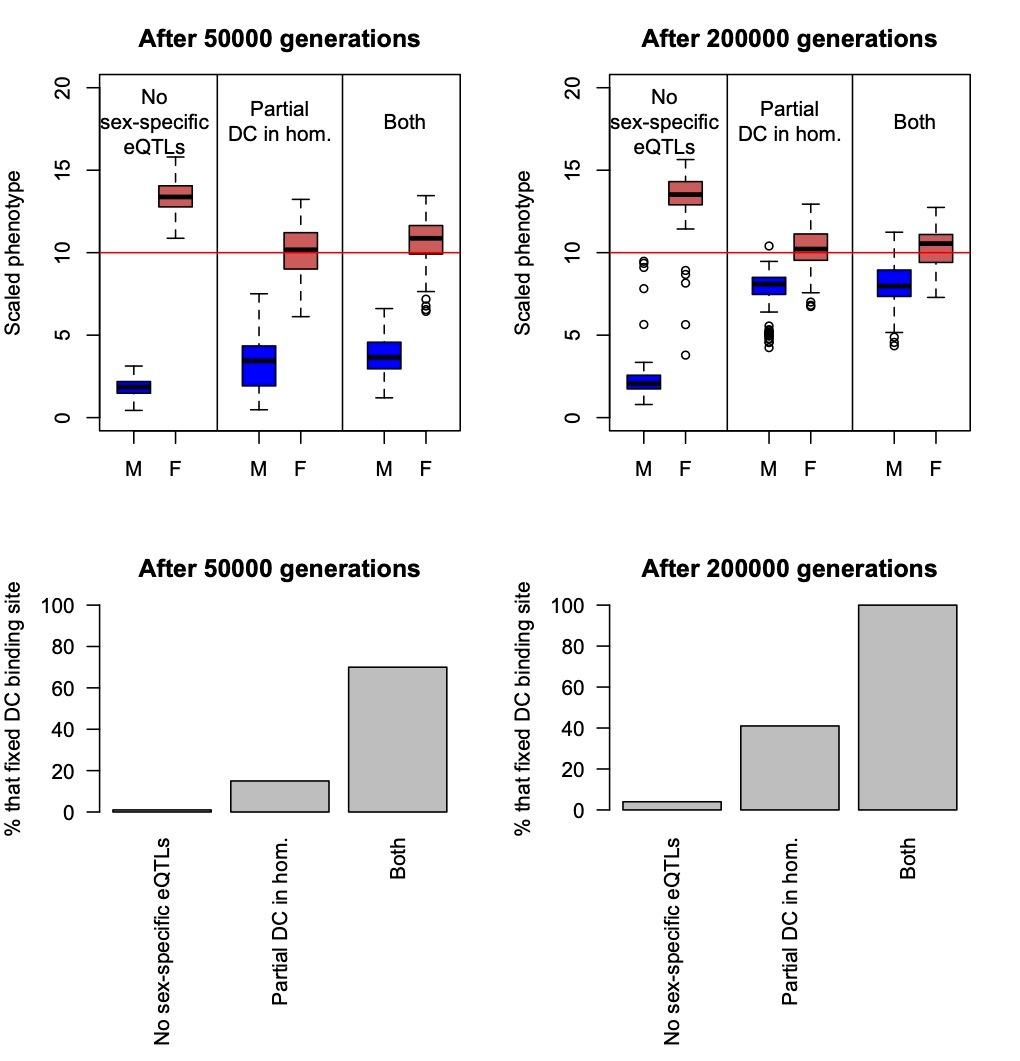

**Supplementary Fig. 15:** Simulations of the evolution of expression of a hemizygous neo-X-linked gene under ancestral dosage compensation involving the down-regulation in the homozygous sex, when no sex-specific eQTLs were allowed, when the downregulation was only partial, and when both conditions were true. Upper panels: Gene expression evolution after 50000 and 20000 generations. The y-axis (Scaled phenotype) represents mean expression across the population, with the red line representing the optimum. M stands for Male and F for Female. Bottom panels: The proportion of simulations of each type where a binding site for dosage compensation was fixed. Each type of simulation was run 100 times.

### **Supplementary Data**

The primer sequences and the location are in this link (<https://seafile.ist.ac.at/f/2c0af2484ed6488189d2/>) and the fasta sequences files used to generate the primer are in this link (<https://seafile.ist.ac.at/f/ce18a09debff46d7ab3f/>).
